# Skip-mers: increasing entropy and sensitivity to detect conserved genic regions with simple cyclic q-grams

**DOI:** 10.1101/179960

**Authors:** Bernardo J. Clavijo, Gonzalo Garcia Accinelli, Luis Yanes, Katie Barr, Jonathan Wright

**Affiliations:** Earlham Institute, Norwich Research Park, Norwich, NR4 7UZ, UK

**Author notes:** Corresponding author: Bernardo J. Clavijo.

## Abstract

Bioinformatic analyses and tools make extensive use of k-mers (fixed contiguous strings of *k* nucleotides) as an informational unit. K-mer analyses are both useful and fast, but are strongly affected by single nucleotide polymorphisms or sequencing errors, effectively hindering direct-analyses of whole regions and decreasing their usability between evolutionary distant samples. Q-grams or spaced seeds, subsequences generated with a pattern of used-and-skipped nucleotides, overcome many of these limitations but introduce larger complexity which hinders their wider adoption.

We introduce a concept of skip-mers, a cyclic pattern of used-and-skipped positions of *k* nucleotides spanning a region of size *S ≥ k*, and show how analyses are improved by using this simple subset of q-grams as a replacement for k-mers. The entropy of skip-mers increases with the larger span, capturing information from more distant positions and increasing the specificity, and uniqueness, of larger span skip-mers within a genome. In addition, skip-mers constructed in cycles of 1 or 2 nucleotides in every 3 (or a multiple of 3) lead to increased sensitivity in the coding regions of genes, by grouping together the more conserved nucleotides of the protein-coding regions.

We implemented a set of tools to count and intersect skip-mers between different datasets, a simple task given that the properties of skip-mers make them a direct substitute for k-mers. We used these tools to show how skip-mers have advantages over k-mers in terms of entropy and increased sensitivity to detect conserved coding sequence, allowing better identification of genic matches between evolutionarily distant species. We then show benefits for multi-genome analyses provided by increased and better correlated coverage of conserved skip-mers across multiple samples.

**Software availability:** the skm-tools implementing the methods described in this manuscript are available under MIT license at http://github.com/bioinfologics/skm-tools/

## 1 INTRODUCTION

Genomes are not random strings, but are the product of millions of years of evolution and selection pressure which imparts unique characteristics to the sequence of nucleotides. These characteristics need to be considered in order to better analyse genomic datasets. Here we exploit the increase in entropy (mean amount of information) from positions that are further away in the genome (Chaisson et al., 2009), and the uneven conservation of coding sequence due to synonymous mutations and the neutral model (Kimura, 1977). The concept of skip-mers extends the familiar concept of k-mers towards a simple cyclic q-gram that can benefit from these two properties.

First, we harness the increased entropy of nucleotides that are further apart by introducing gaps. This has previously been explored to predict regulatory sequences (Ghandi et al., 2014) and to classify sequences taxonomically (Hahn et al., 2016) with q-grams. Instead of general q-gram patterns, we define simple cycles of nucleotide skips which preserve more of the useful properties of k-mers. Second, we take advantage of the increased conservation present in the first two nucleotides of every trinucleotide codon by analysing the skip-mer content of genomes in cycles of three. The consideration of these two concepts allowed us to design skip-mers that improve genic region matches for syntenic analyses.

### 1.1 From k-mers to skip-mers

Bioinformatic analyses make extensive use of k-mers (contiguous strings of k nucleotides) as an informational unit, a concept popularised by short read assemblers (Zerbino and Birney, 2008). Analyses within the k-mer space benefit from a simple formulation of the sampling problem and direct in-hash comparisons (Mapleson et al., 2017). For some analyses, the contiguous nature of k-mers imposes limitations. A single base difference, due to real biological variation or a sequencing error, affects all k-mers crossing that position thus impeding direct analyses by identity. Also, given the strong interdependence of local sequence, contiguous sections capture less information about genome structure and are thus more affected by sequence repetition (Chaisson et al., 2009; Birol et al., 2015).

Q-grams or spaced seeds are strings of nucleotides constructed from a pattern of used-and-skipped positions and have been applied to the sequence matching problem (Kent and Zahler, 2000; Ma et al., 2002; Burkhardt and Kärkkäinen, 2003; Darling et al., 2006). The increased entropy due to a larger span and the higher tolerance to single base differences makes q-grams a better tool than k-mers for many bioinformatics tasks. However, general q-gram analyses can be complicated by the inherent flexibility of the concept and the loss of useful properties of k-mers such as reverse complementability.

We define skip-mers as a cyclic pattern of used-and-skipped positions which achieves increased entropy and tolerance to nucleotide substitution differences by following some simple rules (see Figure 1 and the next section). Skip-mers preserve many of the elegant properties of k-mers such as reverse complementability and existence of a canonical representation which allows strand agnostic analyses (Darling et al., 2006). Also, using cycles of three greatly increases the power of direct intersection between the genomes of different organisms by grouping together the more conserved nucleotides of the protein-coding regions, a property already used by the short 11011011 seeds of the WABA algorithm (Kent and Zahler, 2000). Skip-mers can then be described as a sub-set of q-grams or spaced seeds or a generalisation of the 11011011 seeds first described in WABA: a set of simple cyclic q-grams that increase entropy and sensitivity when analysing divergent coding sequence.

**Figure 1.**
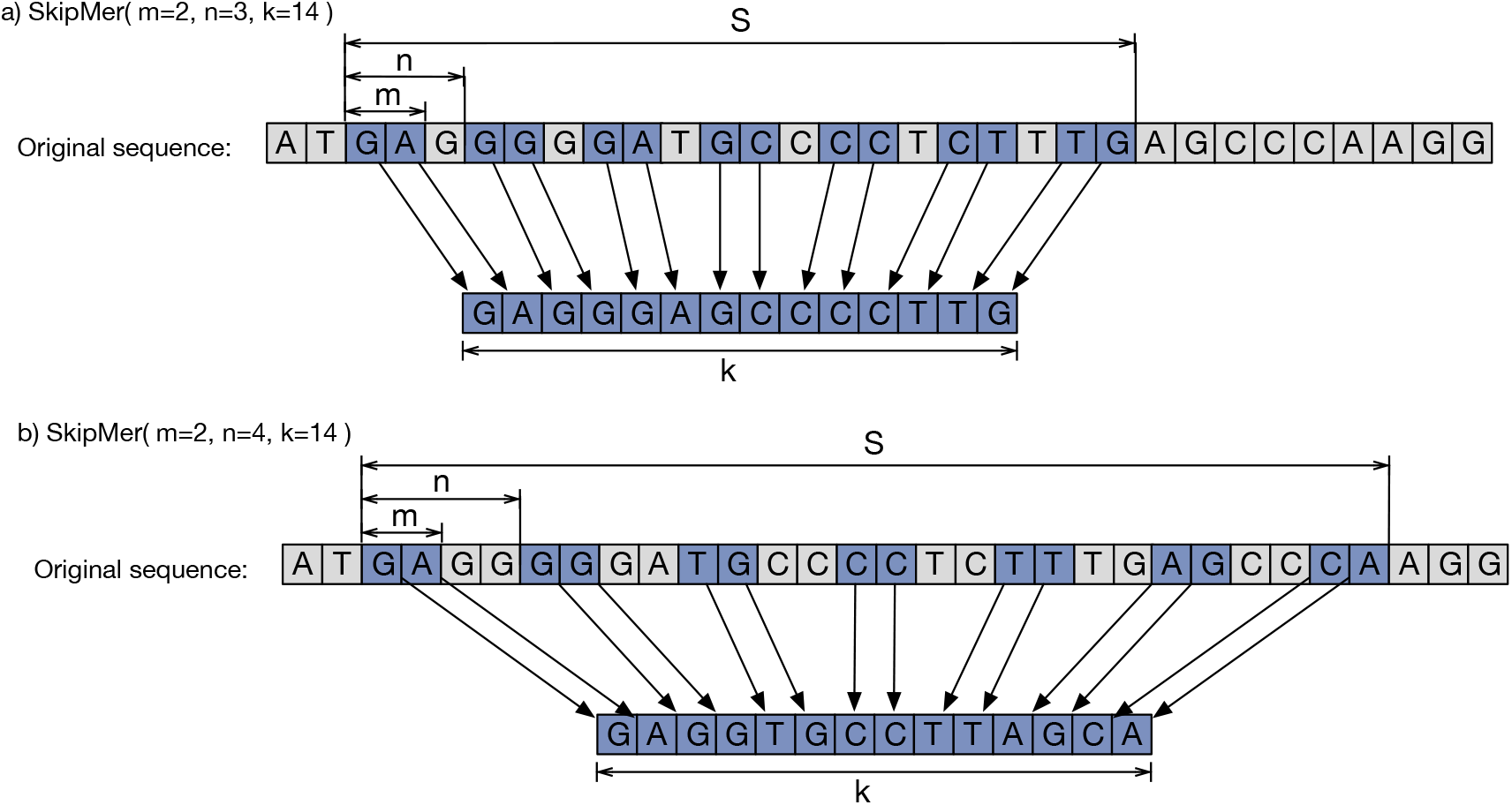
Different *SkipMer(m, n, k)* cycles defined over the same sequence region, resulting in different combinations of bases. The shape of the underlying cyclic q-gram is defined by the variables *m* (used bases per cycle), *n* (cycle length), and *k* (total number of bases).

### 1.2 Skip-mer definition

A skip-mer is a simple cyclic q-gram that includes *m* out of every *n* bases until a total of *k* bases is reached. Its shape is defined by a function *SkipMer(m, n, k)*, as shown in Figure 1. To maintain cyclic properties and the existence of the reverse-complement as a skip-mer defined by the same function, *k* must be a multiple of *m*. This also allows a canonical representation for each skip-mer, defined as the lexicographically smaller of the forward and reverse-complement representations.

Defining *m, n* and *k* fixes a value for *S*, the total span of the skip-mer, given by:

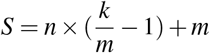

It is important to note that k-mers are a sub-class of skip-mers. A skip-mer with *m = n* will use all contiguous *k* nucleotides, which makes it a k-mer. Throughout this manuscript we often use *m =* 1˄*n* = 1, or the shorter form notations *1-1* or *1-1-k* to refer to k-mers.

## 2 MATERIALS & METHODS

### 2.1 Genome sequences and annotations

To evaluate the properties of skip-mers in a genomic context we used publically available genome assemblies.

Hexaploid bread wheat, *Triticum aestivum* (Clavijo et al., 2017), is a highly repetitive and complex genome, and we used it to investigate the effect of the increased entropy when using larger skip-mer cycles. *Oryza sativa* (Kawahara et al., 2013) and *Brachypodium distachyon* (Vogel et al., 2010) were used as a typical example of synteny in plants, with *Arabidopsis thaliana* (Lamesch et al., 2012) providing a well annotated and distant genome for the 3-way comparisons. *Homo sapiens* (Schneider et al., 2017), *Mus musculus* (Waterston and Pachter, 2002) and *Canis familiaris* (Lindblad-Toh et al., 2005) were used for 2-way and 3-way comparisons between mammal genomes. *Drosophila melanogaster* and eleven other fly genomes (Clark et al., 2007) were used for the multi-way comparison and the presence score analysis. *Escherichia coli* and seven other *Enterobacteriaceae* genomes available from NCBI were used, with accessions: NC_000913.3 (*Escherichia coli*), NZ_CP007557.1 (*Citrobacterfreundii*), NC_003197.1 (*Salmonella enterica*), CP003678.1 (*Enterobacter cloacae*), NZ_CP013990.1 (*Leclercia adecarboxylata*), NC_012917.1 (*Pectobacterium carotovorum*), NZ_CP016889.1 (*Pantoea agglomerans*), andNC_003143.1 (*Yersinia pestis*).

In all analyses where gene regions were used, we downloaded the current GFF3 annotations from Ensembl (Yates et al., 2015) and used a a minimum gene size of 100bp.

### 2.2 skm-tools: skip-mer intersection and coverage analyses

All skip-mer intersection analyses and skip-mer spectra were computed with our skm-tools, available at http://github.com/bioinfologics/skm-tools/. The implementation is based on sorted lists of canonical skip-mers with added attributes such as position on the reference genomes or number of occurrences in a dataset.

In particular, the following tools have been used in the preparation of this manuscript:

**skm-count** counts the number of occurrences of each distinct canonical skip-mer in a fasta input and outputs a spectra histogram.

**skm-multiway-coverage** receives a reference fasta, optionally alongside a GFF3 file and a feature name, and any number of extra datasets. The intersection of skip-mers from all the extra datasets is computed versus the reference dataset, shared-by-all skip-mers statistics are reported as each genome is processed (See Figure 4 for an example progression). If a GFF3 and a feature name is provided, the output will classify the skip-mers according to their presence in regions annotated with the feature, and a file with details of coverage for each feature by each of the extra datasets will be produced.

The current implementation of the skm-multiway-coverage tool includes a coverage cut-off that defaults to 1 as this is appropriate for the current study. All skip-mers that are at a higher frequency than the cut-off are eliminated before any analysis. To consider candidate matches for alignment of conserved sequence it is appropriate to discard skip-mers with a higher copy number than your expected number of matches as this will filter repetitive matches including background noise. While our current choice of cut-off at 1 makes sense in a general analysis as the one presented in this manuscript, care needs to be taken to make reasonable choices for future applications.

### 2.3 Coverage score

The coverage score is used as a proxy for sequence conservation. To approximate a measure of conserved nucleotides, the coverage is projected over individual nucleotides rather than directly counting shared skip-mers which would introduce redundancy from phased matches. An equivalent coverage metric for spaced seeds can be found in Noé and Martin (2014) where it is also used to estimate distances. The score for each feature (i.e. gene) versus each genome in the multi-way analyses is calculated as the total number of bases that are included in matching skip-mers from that genome divided by the total number of bases that are covered by valid (i.e. copy number below threshold in the reference) skip-mers from the reference:

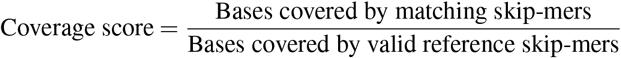

The coverage cut-off is applied before any analyses are performed. When using the default cut-off of 1, skip-mers that have a higher copy number in the reference will not be evaluated for scoring and skip-mers that have a single copy in the reference but more than one copy in the scoring genome will not be counted as covered.

## 3 RESULTS

### 3.1 Increasing a skip-mer cycle length and span increases specificity

We analysed a genome assembly of *Triticum aestivum* to investigate the effect of the cycle size *n* in the multiplicity of the skip-mers in a genome. Figure 2 shows how increasing *n*, and thus the total span of a skip-mer (*S*), increases the entropy for each skip-mer. The increased entropy decreases the number of copies of each distinct skip-mer in the genome. This ultimately results in more unique skip-mers. In the wheat genome, there are more than twice as many unique skip-mers using *SkipMer*(1,16,31) as there are using *SkipMer*(1,1,31) which corresponds to a 31-mers.

**Figure 2.**
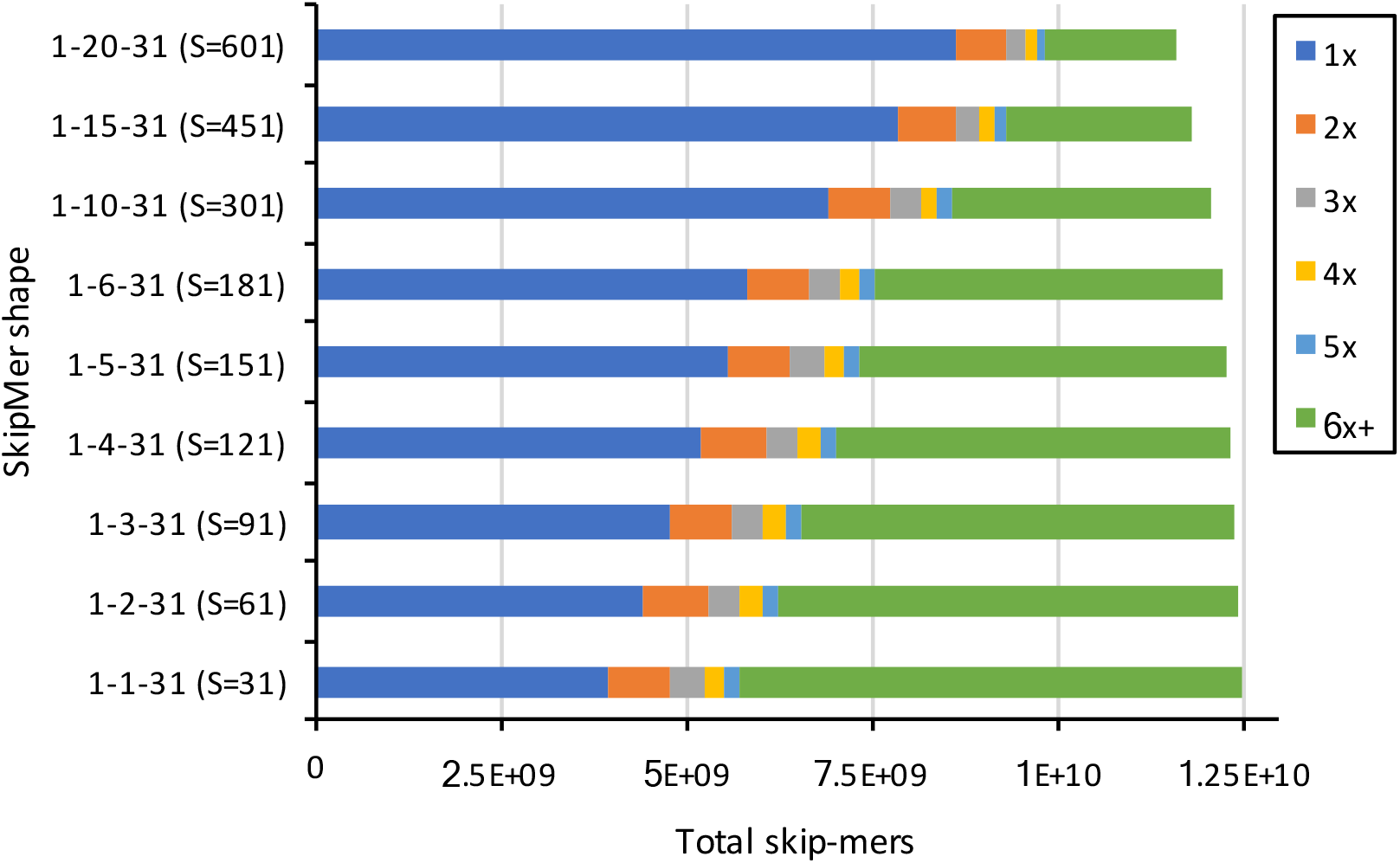
Multiplicity as a function of *n* for skip-mers in *Triticum aestivum*. Each bar represents the total number of skip-mers in the assembly, so a skip-mer appearing *x* times contributes *x* units in the *x* component. All skip-mers use 31bp (*k* = 31) and 1bp per cycle (*m* = 1).

### 3.2 Using triplet-based cycles increases perfect skip-mer matches in conserved genic sequence between species

Synonymous mutations which are not removed by purifying selection, because they do not affect the amino-acid encoded by a trinucleotide codon, produce a cycle-3 modulation in conserved coding regions (Kimura, 1977). Skip-mers with cycle lengths that are a multiple of 3 (*n* = 3*c*) group first and/or second nucleotides in subsequent in-frame codons, to increase sensitivity on *SkipMer*(*m,n* = 3*c,k*) to detect conserved coding regions.

Figure 3 shows how, for the 2-way intersections in (a) and (c), the shared skip-mers in non-genic regions decrease as the span increases, in agreement with the increase of entropy and thus uniqueness. In genic regions, this higher entropy is combined with increased sensitivity for coding sequence resulting in increased matches when *n* = 3*c*. This effect is further accentuated in the 3-way intersections in (b) and (d), due to independent synonymous mutations in the 3 genomes.

**Figure 3.**
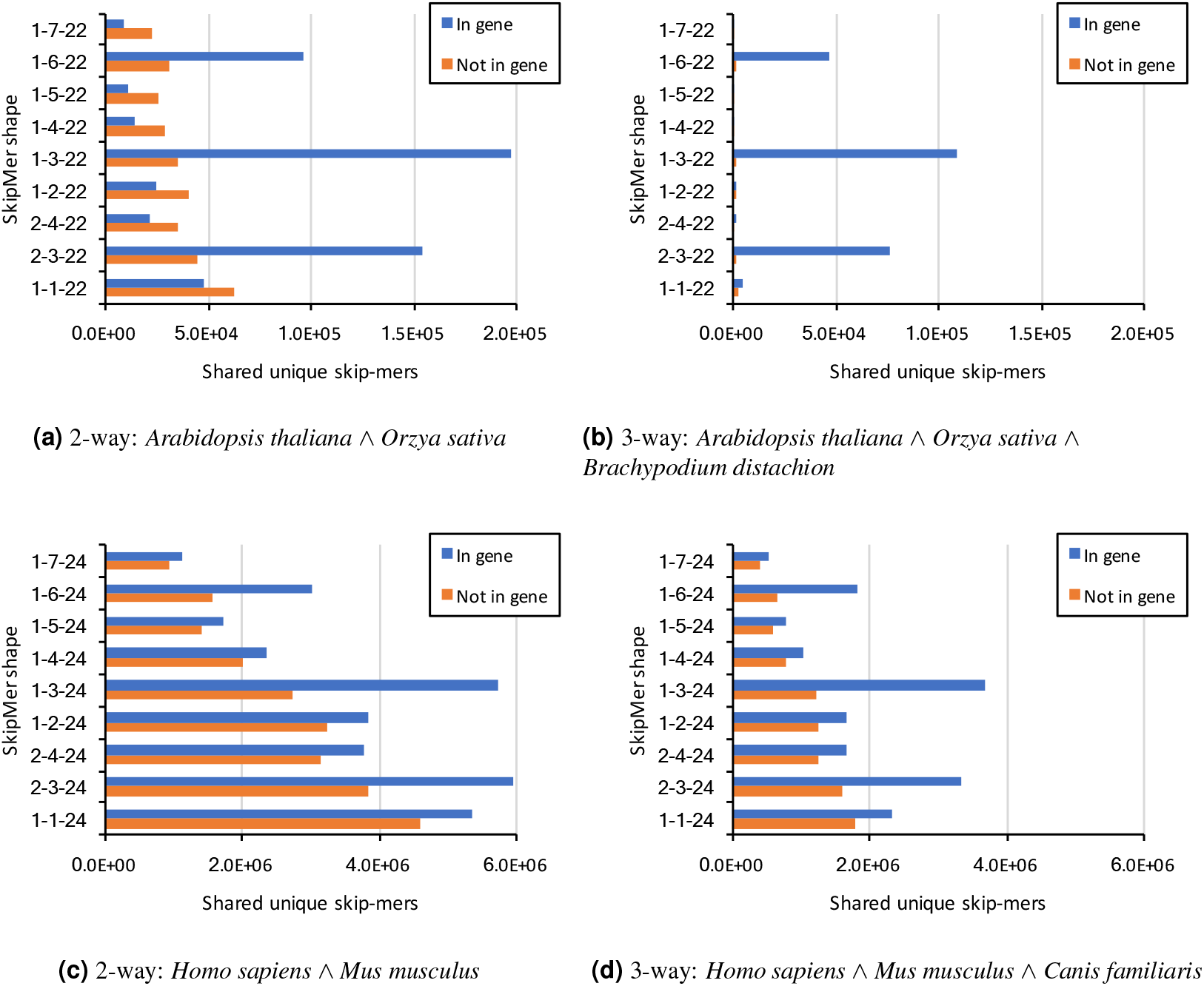
Effect of different combinations of *m* and *n*, while keeping *k* constant, for 2-way and 3-way skip-mer intersections. Only unique skip-mers are considered and skip-mers originating from sequence annotated with gene features on the first genome are classified as *“In gene”*. The skip-mer shapes are sorted along the vertical axis according to total skip-mer span (*S*), with the largest span on top.

The best result in terms of sensitivity for three of the four examples is produced by *SkipMeri* 1.3. *k)* which groups the nucleotides according to their position in the codon. While *SkipMer*(2,3,24) presents a slightly larger number of “In gene” matches in (c), *SkipMer*(1,3,24) with its larger span decreases the number of ‘‘Not in gene” matches, which makes it a better choice.

### 3.3 Conserved sequence from n=3 skip-mers enables direct intersection analyses across many samples at diverse evolutionary distances

One of the limitations of k-mers for direct intersection analyses among many samples is the decrease in probability of finding kmers that are shared across all of the samples. The results from our 2-way and 3-way analyses show that skip-mers are more sensitive to conserved coding sequence. We intersected both the twelve *Drosophila* genomes from Clark et al. (2007) and the *Enterobacteriaceae* dataset to explore how coverage over the reference from unique shared skip-mers decreases both in genic and non-genic regions as we progressively include more samples.

In Figure 4 (a) the eleven other genomes are incorporated into the *Drosophila melanogaster* based analysis starting from the closest to *D. melanogaster* in the phylogeny proposed by Clark et al. (2007): *D. simulans, D. sechellia, D. yakuba, D. ananassae, D. erecta, D. pseudoobscura, D. willistoni, D. virilis, D. mojavensis*, and *D. grimshawi*. In Figure 4 (b) this order is reversed, starting from *D. grimshawi* and ending with *D. simulans*. The order of *Enterobacteriaceae* genomes in Figure 4 (c) is: *Citrobacter freundii, Salmonella enterica, Enterobacter cloacae, Leclercia adecarboxylata, Pectobacterium carotovorum, Pantoea agglomerans*, and *Yersinia pestis*. In Figure 4 (d) this order is reversed, starting from *Yersinia pestis* and ending with *Citrobacter freundii*.

**Figure 4.**
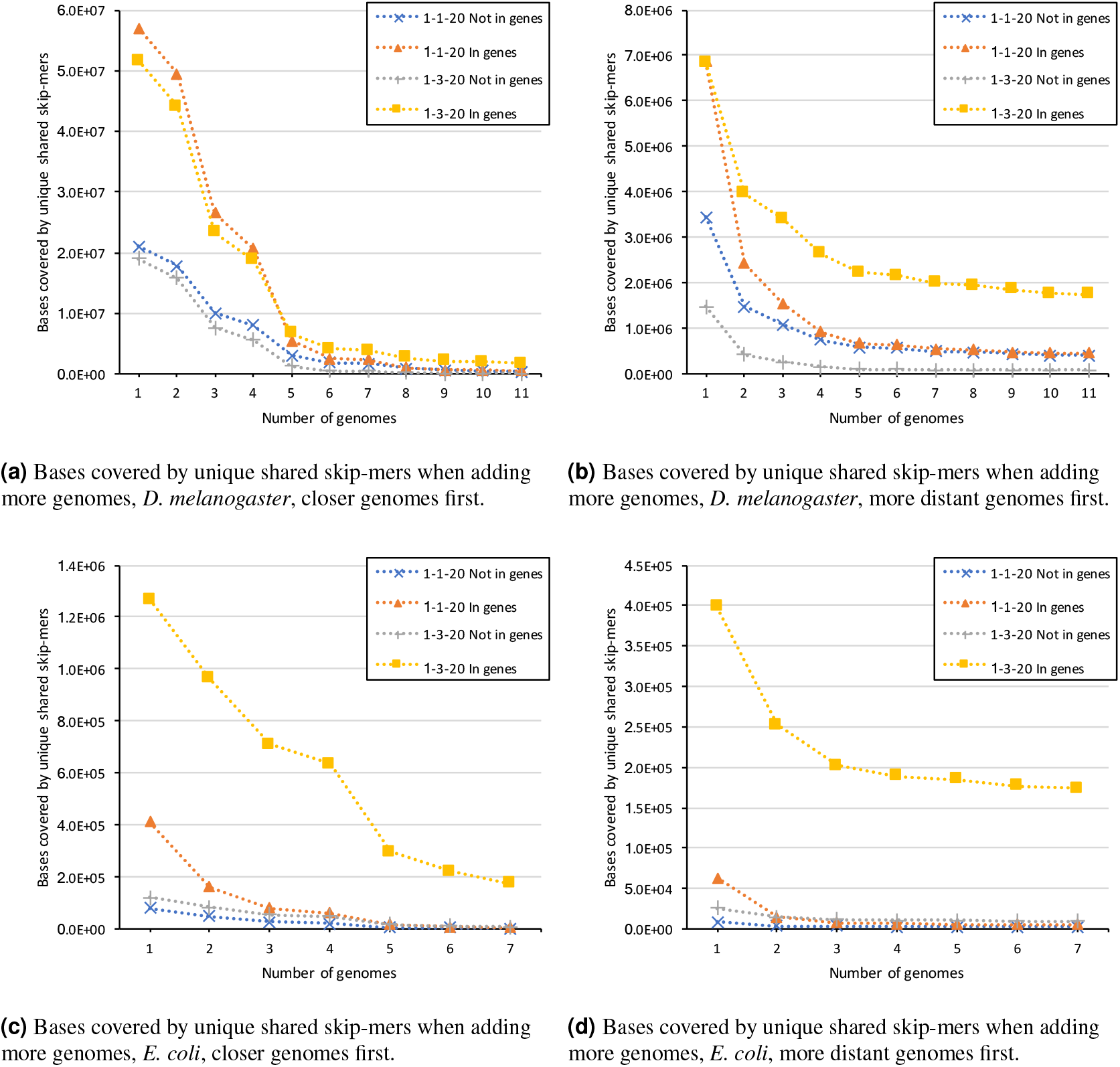
Bases covered by unique shared skip-mers shared across all genomes, for sets of different numbers of genomes from the 12 *Drosophila* dataset and the 7 *Enterobacteriaceae* genomes.

In every analysis shown in Figure 4 the genic intersection computed by *SkipMer*(1,3,20) is less affected by the introduction of extra genomes due to the increased sensitivity in the conserved coding regions. In Figure 4 (a), the first four genomes show a small increase in sensitivity when using k-mers, due to their closeness to the reference, but the effect is reversed after the incorporation of the fifth genome. In Figure 4 (b), the first genome shows only a difference for “not-in-gene” matches, with the cycle-3 skip-mers being less conserved outside coding constraints; from the second genome onwards, the advantages of skip-mers become more evident. In Figures 4 (c) and 4 (d) the larger evolutionary distance increases the effects of skip-mer conservation in the analysis. All these results show how skip-mers can be used to provide a small set of conserved sequences across the diverse genomes in the datasets.

### 3.4 Coverage of matching sequence across many samples using skip-mers with n=3 shows higher correlation than using k-mers

To explore the advantages of the *SkipMer*(1,3, *k*) analyses across divergent genomes we compared the properties of sequence coverage for *Drosophila melanogaster* to the eleven other *Drosophila* genomes and for *Escherichia coli* to the seven other *Enterobacteriaceae* using *SkipMer*(1,1,20), which is equivalent to a 20-mer, and *SkipMer*(1,3,20). We implement a base coverage score as described in section 2.3 and assigned each gene with a length of 100bp or more in the reference a coverage score between 0 and 1 for each of the genomes.

A distribution analysis for the scores per genome (Supplementary Figure S1) shows the more divergent genomes increase their scores for sequence coverage in genes when using *SkipMer*(1,3,20). This reflects the increased sensitivity of cycle-3 skip-mers within coding regions. This score increase is particularly large in the bacteria dataset, where the samples are more evolutionary distant.

We computed correlations between the gene scores for *D. melanogaster* from every pair of the other 11 genomes for both *SkipMer*(1,1,20) and *SkipMer*(1,3,20) (See Supplementary Material Tables SI and S2, and Figures S2 and S3). Figure 5 shows the comparison between each genome-pair correlation on *SkipMer*(1,1,20) and *SkipMer*(1,3,20). There is increased correlation when using cycle-3 skip-mers, with larger relative improvements on the less correlated genome pairs. This suggests cycle-3 skip-mers provide a more robust coverage score which can be better used as a proxy for evolutionary pressure and selection.

**Figure 5.**
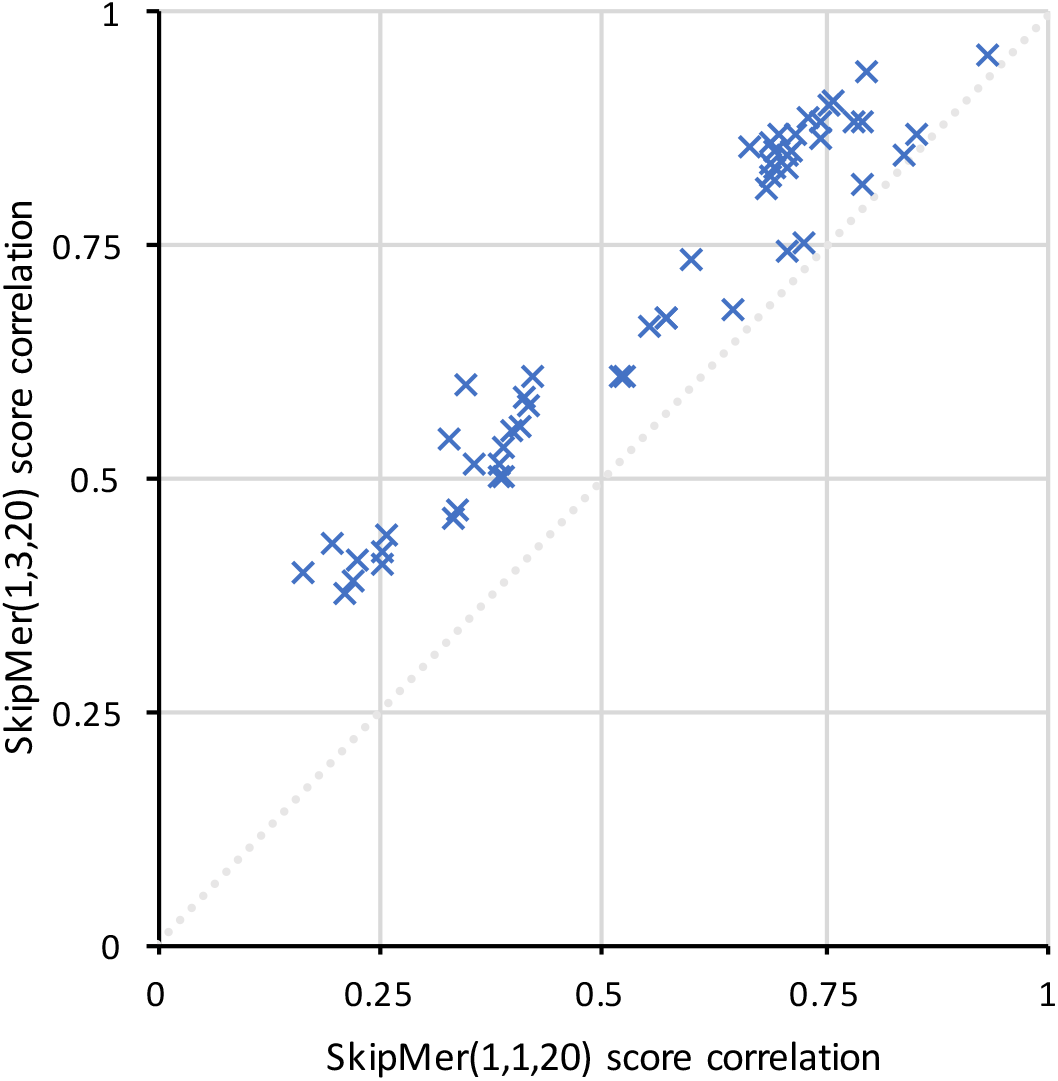
Comparison of correlations for the coverage scores of the *Drosophila melanogaster*. Points above the 1:1 line represent genome pairs with better score correlation in *SkipMer*(1,3,20) than in *SkipMer*(1,1,20) (equivalent to 20-mers).

The score for the *Enterobacteriaceae* dataset when using k-mers is so low, and the effect of gene divergence and gain/loss so pronounced, that computing a correlation for scores would not correspond with conservation differences without a more elaborate analysis. The scatter plots of scores in Supplementary Figures S4 and S5 illustrate this point and the improvements on scores with skip-mers.

## 4 DISCUSSION

Increasing the span of skip-mers increases their entropy when sampled from a genome. Using this increased-entropy analysis unit rather than k-mers will enable more informative analyses with small adaptations to existing techniques. We expect this key feature of the data points having the same amount of data (bp) but increased entropy to enable more exhaustive or significant analyses in roughly similar computational space and time.

The analysis of sequence in cyclic groups of *n =* 3 increases sensitivity to detect conserved coding sequence by grouping the nucleotides in synchronisation with the codon positions. In the typical case of *SkipMer*(*m* = 1, *n* = 3, *k*) there will be, for the same group of *k* contiguous codons, a skip-mer containing all first nucleotides, a skip-mer containing all second nucleotides and a skip-mer containing all third nucleotides. This grouping increases perfect matches in genes from first-nucleotide position and second-nucleotide position skip-mers, providing an alternative to the use of protein translation analyses.

When computing multi-genome intersections, there is a stronger signal of conservation across multiple divergent genomes from *n* = 3 skip-mers than from contiguous sequences such as k-mers. Because of the better correspondence between sequence that is actively conserved and the set of matches, the reduction in matches due to the addition of extra genomes in intermediate positions in the phylogeny is less pronounced. Also, the coding sequence coverage by direct matches is a more robust metric, which enables the direct comparison of results produced from different sets of genomes.

A complementary effect to the concentration of more conserved sequence, from first and second nucleotides, in the cycle-3 skip-mers is the concentration of more variable sequence, from third nucleotides, in a small number of skip-mers. In the preceding analyses, these more variable nucleotides have been discarded with the noise and repetitions. For applications where a weak signal for variation needs to be analysed, skip-mers can be leveraged to provide a very high entropy set of sequences to give increased discrimination power.

Our results, in addition to confirming and expanding previous work on q-grams and spaced-seeds, suggest skip-mers will have a wide range of applications in bioinformatic analyses. For whole-genome and multi-genome alignment, skip-mers will provide accurate conserved seeds, and more specific matches in complex regions. For evolutionary analyses, skip-mers will allow improved detection of functionally equivalent regions. For RNA-seq and exome analyses, skip-mers will provide a meaningful set of starting seeds or a projection base, thus enabling more distant samples to be analysed together either against a reference or in a reference-free manner.

Skip-mers will also be useful in raw read analyses. For classification of sequences, or species detection, skip-mers will provide better clustering of coding regions from a common origin, and could even be used to estimate conservation scores for single reads. Aligning skip-mers from raw reads to one or many references will guide the reconstruction of conserved regions while considering novel variants. These conserved region intersected representations can then be used to quickly characterise the genic space of a genome.

## CONCLUSIONS

We have shown how skip-mer based analyses benefit from extra entropy and sensitivity to outperform k-mer based analyses given the non-random nature of genomic sequence. These principles stand across a wide range of prokaryotic and eukaryotic genomes and in different multi-genome scenarios, improving the analysis of conserved coding regions. Common k-mer based techniques can easily adopt skip-mers, due to their many shared properties. Both constructions are reversible strings of nucleotides that can be made strand-agnostic with canonical representations. In general, with a genomic landscape that is shifting to in-field sampling and exploring more diversity than ever before, we expect skip-mers and other evolution-friendly information units to provide the basis for a new generation of biological analyses.

## AUTHORS CONTRIBUTIONS

BJC and GG developed the initial concepts and discussed implications and refinements over time. BJC implemented the first version of the *skm-tools*, ran the analyses and produced the first draft of the manuscript. LY contributed optimisations and improvements for the *skm-tools*. All authors tested the *skm-tools*; discussed analyses, results and improvements; and contributed to the final version of the manuscript.

## ACKNOWLEDGMENTS

The authors would like to thank Wilfried Haerty for his support on the analysis of the *Drosophila* genomes, and C. Titus Brown for his input on the microbial genomes analysis. The authors also thank Erik Garrison, Mark McMullan, Neil Hall, Norma Paniego, Manfred Grabherr, Laurent Noé and Federica di Palma for their useful comments and feedback.

